# Circuit-specific reorganization of hippocampal-entorhinal-prefrontal subnetworks supports memory recall across the lifespan

**DOI:** 10.64898/2026.08.10.743925

**Authors:** Erika Atucha, Rukhshona Kayumova, Celia Fürst, Magdalena Sauvage

## Abstract

How memories reorganize across brain circuits as they age remains a central question in systems neuroscience. Systems consolidation is thought to progressively shift memory reliance from the hippocampus to distributed cortical networks, yet the contribution of cortical regions beyond the prefrontal cortex and the nature of this shift remains unclear. Here we define the circuit-level organization of remote memory recall across entorhinal, prefrontal, and hippocampal subregions. Using high-resolution activity mapping combined with causal manipulations that leverage natural memory decay, we adapted a murine object-location paradigm to examine memory recall across the lifespan. We find that recall of early remote memories (1 month) selectively depends on a LEC-hippocampal (CA1/CA3) circuit, whereas recall of older memories (6-12 months) recruits a distinct and broader network involving both LEC and MEC together with ACC and CA1. These findings reveal a temporally ordered, circuit-specific reconfiguration of hippocampo-cortical networks and identify the EC as a dynamic hub in remote memory retrieval. Our results refine prevailing systems consolidation theories by showing that memory consolidation is a circuit-specific and temporally ordered process, rather than a passive gradual phenomenon, and position the EC as a central and dynamic component of remote memory retrieval alongside the PFC.

**Graphical abstract:** 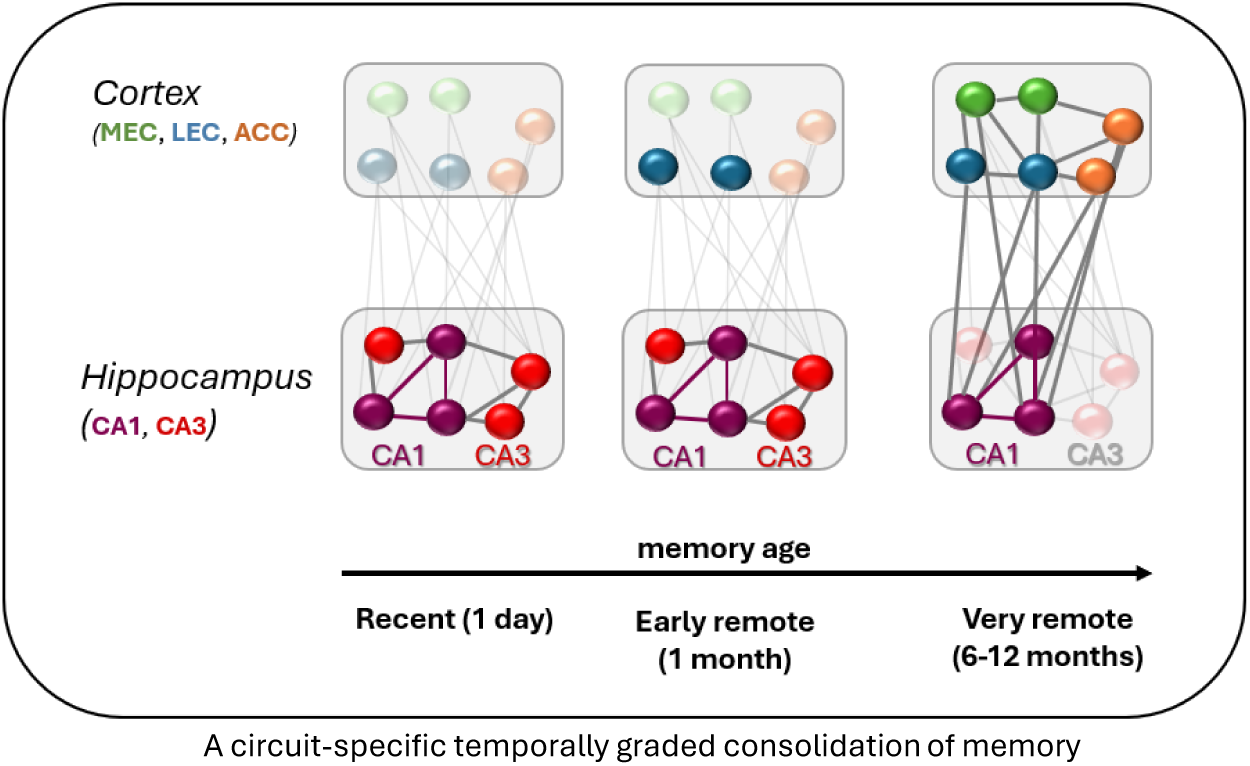

## Introduction

How to recall unique events such as the memory of one’s birthday, is a defining feature of episodic memory and a major focus in memory research. Extensive work in humans and animals has identified the hippocampus (HIP) and the prefrontal cortex (PFC) as core components of the neural circuitry supporting memory consolidation and retrieval over time ^1–7^. Both structures comprise anatomically and functionally distinct subregions - such as CA1 and CA3 within the HIP ^8–12^, and the anterior cingulate (ACC), prelimbic (PL), and infralimbic (IL) cortices within the PFC ^13–15^ whose specific contributions to remote memory recall over time remain poorly understood. Progress in resolving these contributions has been limited, at least in part, by methodological constraints. Indeed, in humans, standard neuroimaging approaches, such as 3T fMRI, lack the spatial resolution required to dissociate activity in adjacent subregions, making it difficult to identify the precise neural substrates of memory recall. This limitation is particularly relevant in light of the prevailing view that memories become increasingly distributed across cortical regions as they age ^3,7^. Moreover, although this framework is supported by substantial evidence, research on memory consolidation has focused primarily on the PFC, leaving the involvement of other cortical regions largely unexplored. Yet, recent findings have implicated the entorhinal cortex (EC) - the main cortical input to the hippocampus - as a candidate contributor to memory recall. The EC contains functionally distinct lateral (LEC) and medial (MEC) subregions, which differentially process nonspatial and spatial information ^11,16–22^. In addition, emerging evidence suggests that EC subregions become increasingly engaged during the recall of remote memories and provide targeted support to the hippocampus, particularly the CA1 subfield ^23,24^. However, whether the EC subregions are necessary for remote memory recall, how they specifically contribute across memory age, and whether these dynamics unfold over timescales relevant to human memory remain unclear.

Addressing these questions is complicated by major differences between human and animal studies. Rodent work on systems consolidation has primarily relied on fear-based paradigms, whereas human studies typically examine more neutral memories, such as words or objects, over a much wider time-window (half a lifetime in humans compared to a month in rodents). Moreover, neither approach has yet combined behaviorally relevant tasks with the spatial resolution needed to dissociate activity across entorhinal and prefrontal subregions.

To investigate whether the consolidation of memory over time reflect a passive time-dependent process or a circuit-specific reorganization bound to memory demands, here, we identified the specific contributions of entorhinal (LEC, MEC), prefrontal (ACC, PL, IL), and hippocampal (CA1, CA3) subregions to memory recall across time using experimental conditions designed to more closely align animal and human memory research and to yield cellular spatial resolution. To do so, we adapted a murine object-location task to examine the neural substrates of memory recall across an extended temporal window, ranging from recent (1 day and 1 week) to early remote (1 month) and very remote (6 months and 1 year) time points (Fig. 1A; 1 year corresponding approximately to half the lifespan of a mouse, which is comparable to the time window commonly used in human memory research). To establish a causal relationship between neural engagement and memory performance, we leveraged the natural decay of memory over time and compared memory-intact and memory-impaired mice at identical memory ages. Neural activity at test was mapped with high resolution in entorhinal (LEC, MEC), prefrontal (ACC, PL, IL) and hippocampal (CA1, CA3) subregions, by detecting the RNA of the immediate-early gene *Arc* (Fig. 1A), whose expression is closely tied to synaptic plasticity, reflects memory demands and is commonly used to mapping mnemonic circuits in rodents^25–31^.

**Fig. 1.**
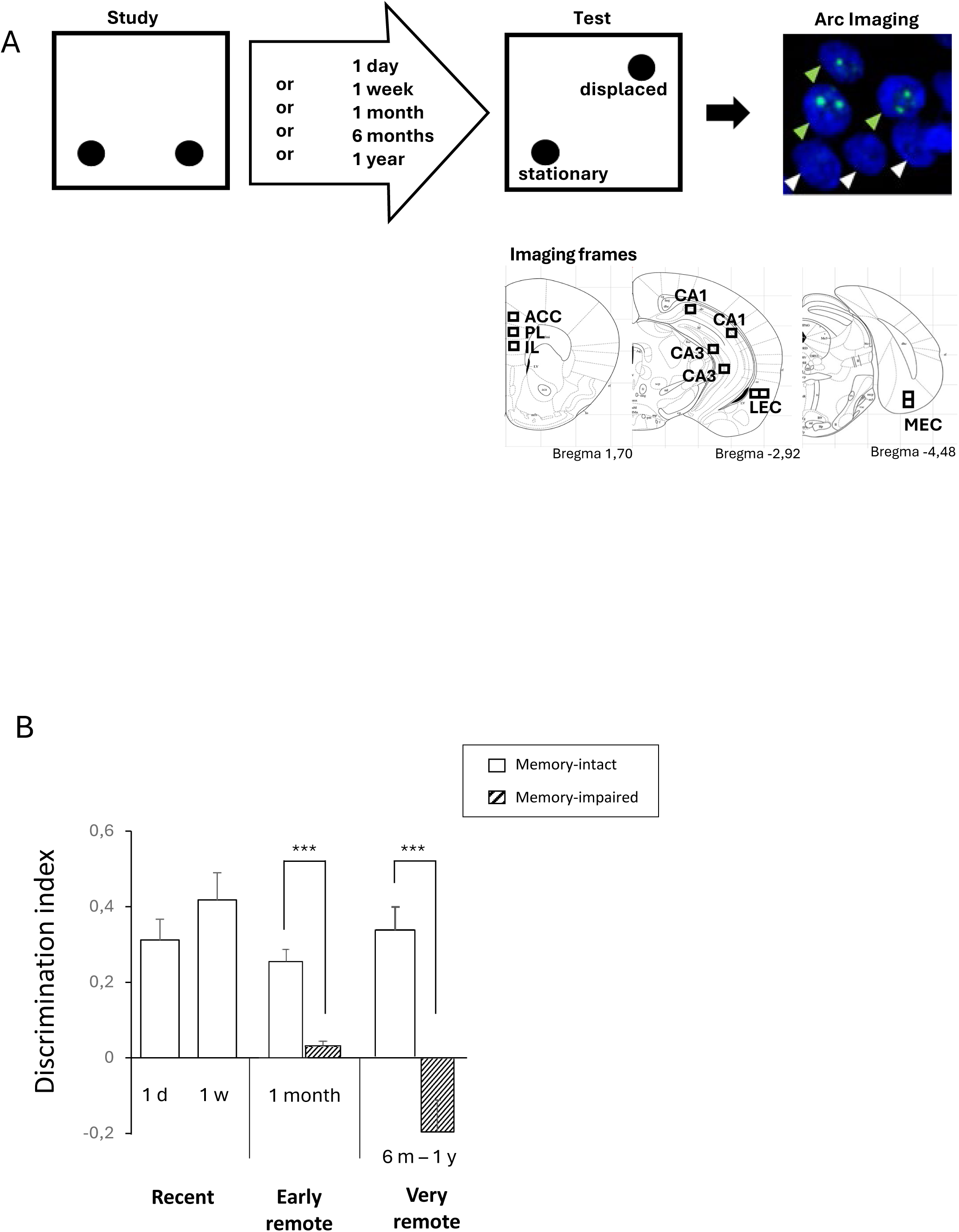
Behavioral task, imaging frames and memory performance. **A)** *Behavioral protocol.* Animals were placed in an open field containing two identical objects during a study phase and then returned to their home cage. After a delay (1 day, 1 week, 1 month, 6 months or 1 year), animals were re-exposed to the same open field with two copies of the original objects, with one object displaced to a novel location. Following the test, mice were sacrificed and brains were processed for the detection of pre-mRNA of the immediate early gene *Arc* (green arrows, examples of CA1 *Arc*+ cells; white arrows, *Arc*-cells; nuclei labeled with DAPI, blue). *Imaging frames*. Black frames indicate the levels at which images were acquired using a 40× objective. Three images were collected per target area from equidistant, non-consecutive sections. Cell counting was restricted to neurons, as described in Vazdarjanova & Guzowski^61^. **B)** *Memory performance*. Discrimination indices (DI) were calculated as the difference between time spent exploring the displaced versus stationary object, divided by total object exploration time. Memory-intact animals (white bars) exhibited positive DI values significantly above chance level (one-sample t-test against 0, all Ps < 0.05), indicating successful memory recall. Memory-impaired animals (striped bars) showed DI values comparable to zero (Ps > 0.06), indicating impaired recall of the original object location. All mice successfully recalled object locations at recent time-points (1 day and 1 week), whereas only half of the populations did so at more remote time points (1 month and 6-12 months). ***P < 0.001. Bars represent mean + SEM.

We found that, although recalling recent object-location memories relies predominantly on CA1 and CA3, LEC is additionally selectively required for early remote memory recall. In contrast, recalling even more remote memories depends on the functional integrity of both LEC and MEC and selectively on the ACC and CA1 among the PFC and hippocampal subregions. These results suggest a circuit-specific and time-dependent network reconfiguration for memory recall rather than a passive cortical distribution of the memory traces over time and identify the EC as a dynamic hub in remote memory recall alongside the PFC.

## Results

### Performance in mice with intact memory is enhanced compared to memory-impaired mice

One of the aims of the present study is to further bridge human and animal memory research. To this end, we chose to study neutral memories (memory for the location of objects) and adapted a standard murine protocol for the detection of memory traces as old as those studied in humans (20-40-year-old memories, roughly corresponding to 6- and 12-month-old memories in mice based on life expectancy). A total of n= 26 mice were tested using this adapted version of the object-location task. Memory performance was evaluated in distinct groups of mice 1 day, 1 week, 1 month, 6 months and 12 months after memory formation (i.e., after the study phase; Fig. 1A). According to the novelty-preference principle ^32^, successful recall of the original location of the objects is reflected by a longer exploration of the displaced object compared to the stationary object at test, yielding a positive discrimination index (DI > 0; see Material and Methods for DI calculation). Mice were classified into memory-intact and memory-impaired groups using K-means clustering based on their DI^33^ (see Materials & Methods). Successful memory recall was observed at the recent time points for all mice (1 day and 1 week) and, in agreement with the literature ^23,24^, in half of the populations one and six months after memory formation (one-sample comparisons to 0: memory-intact groups, *Ps* < 0.05; Fig. 1B). In addition, memory performance across memory-intact groups was comparable up to 6 months after encoding (one-way ANOVA, memory-intact groups: F(3,12) = 1.45; *P* = 0.277; Fig. 1B), indicating that the strength of the memory recalled was comparable over half a year across memory-intact groups.

Furthermore, failure to recall was observed in the remaining halves of the one-and six-months old memory groups in line with the literature ^23,24,34^ as well as in all mice one year after memory formation (of note, mice with impaired memory 6- and 12-months after study were pooled into a single 6 month-1 year memory-impaired group as their DIs did not cluster according to the age of the memory (K-means clustering) and groups show similar mean DIs (*P* = 0.45; one-sample comparisons to 0: memory-impaired groups, 1 month or 6m-1y: *Ps* > 0.05). Finally, comparisons of the discrimination indices between memory-intact and memory-impaired groups revealed higher performance in memory-intact than in memory-impaired groups when early (1-month) or very remote (6- or 12-month old) memories were recalled (two-way ANOVA, performance × delay interaction effect: F(1,14) = 5.48, *P* = 0.034, main performance effect: F(1,14) = 33.82, *P* < 0.001; no main delay effect: *P* = 0.268; post hoc comparisons, memory-intact vs impaired: early remote: *P* = 0.001; very remote: *P* = 0.001; Bonferroni-Holm corrected. These comparisons were not performed at the recent time points (1 day and 1 week) because of the absence of memory-impaired mice at these delays).

Altogether, these results indicate that mice with intact memory show comparable performance for up to half a year and reveal the emergence of a memory-impaired subpopulation when memories reach one month of age. This subpopulation comprises approximately half of the group and ultimately expands to encompass the entire population after one year. Next, we compared neural activity in entorhinal cortex (EC), prefrontal cortex (PFC), and hippocampal (HIP) subregions between memory-intact and memory-impaired mice to determine which regions are necessary for recalling object locations.

### The LEC is necessary for the recall of younger memories than the MEC

Given the recent reports indicating a potentially important role of the EC subareas in system memory consolidation, we first investigated whether the LEC and the MEC play an essential role in supporting the recall of object-location memory within this framework. To do so, we first assessed the extent of their engagement during successful memory recall by evaluating the task-induced proportions of *Arc*+ cells in memory-intact mice (see Materials & Methods) and subsequently compared those proportions to those of memory-impaired mice. In memory-intact groups, the LEC and MEC were recruited independently of the age of the memory during recall, and proportion of task-induced *Arc*+ cells were maximal at remote time points (blue and green bars, respectively; Fig. 2A; comparisons to 0: all *Ps* < 0.015; one-way ANOVAs, LEC: F(3,12) = 59.25, *P* < 0.001 and MEC: F(3,12) = 12.45, *P* < 0.001; Tukey post hoc tests: early or very remote time-points vs recent time-points: *Ps* < 0.001; 1-day vs 1-week or 1-month vs 6-12 months (*Ps* > 0.65), Bonferroni-Holm corrected). In addition, *Arc* expression in memory-intact groups did not differ between LEC and MEC over time (two-way ANOVA, delay × region interaction effect: F(3,12) = 2.27; *P =* 0.132; other effects: *Ps* < 0.05). Hence, these results indicate a similar recruitment of LEC and MEC during successful memory recall over time with a maximal engagement for remote memory recall.

**Fig. 2.**
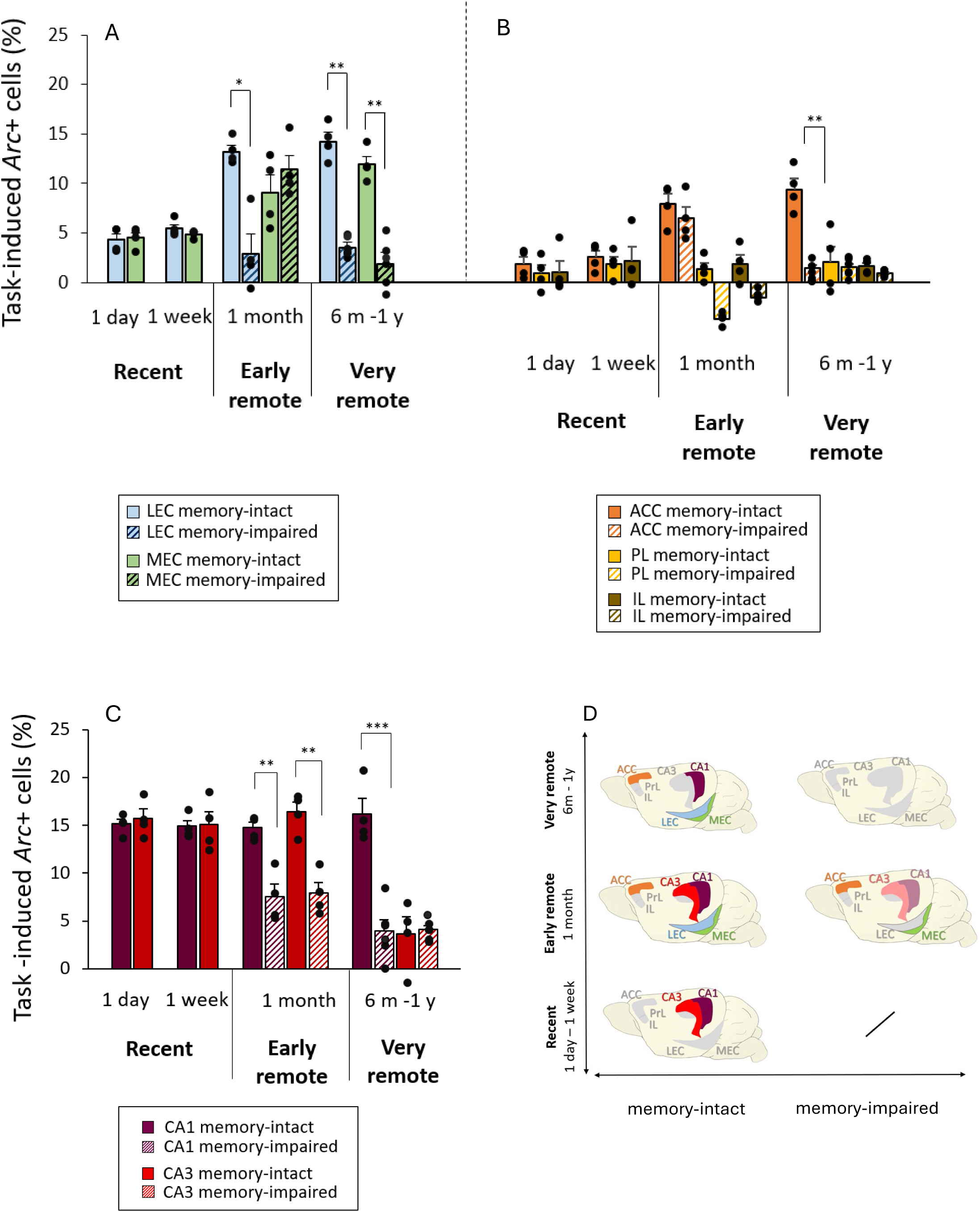
Neural activity at test in entorhinal, prefrontal and hippocampal subareas of memory-intact and memory-impaired mice over time. Task-induced activity was assessed in the LEC, MEC, ACC, PL, IL, CA1 and CA3 at recent (1 day and 1 week), early remote (1 month) and very remote (6-12 months) time points by detecting pre-mRNA of the immediate-early gene *Arc* using *in-situ* hybridization ^30,31,61^. **A and B).** All cortical regions (LEC, MEC, ACC) were maximally recruited at the remote delays (memory-intact mice: one-way ANOVAs, Ps < 0.001; Tukey’s post hoc tests early or very remote delays vs recent delays: *P*s< 0.05). Comparisons of task-induced *Arc* expression between memory-intact and memory-impaired mice shows that, at 1 month, only LEC activity was significantly reduced in impaired animals, indicating a predominant role for LEC over the MEC and ACC in early remote object-location memory recall. In contrast, activity levels were lower in all three regions in mice with deficits when memories were older, suggesting increasing contributions of MEC and ACC in the recall of this type of memory as they age (three-way ANOVA, all effects P< 0.05; posthoc comparisons memory-intact vs memory-impaired mice: 1 month: LEC P = 0.007, MEC and ACC: Ps = 0.721; 6-12 months: MEC, LEC, ACC all Ps < 0.001, Bonferroni-Holm corrected). **C)** In the hippocampus, CA1 was recruited at test irrespective of memory age, whereas CA3 was no longer engaged for the recall of the most remote memories (memory-intact mice: one-sample t-tests against 0: CA1, all Ps < 0.01; CA3: 1 day, 1 week, 1 month: Ps < 0.01, 6-12 months P = 0.132; Bonferroni-Holm corrected). Comparisons of task-induced *Arc* expression between memory-intact and memory-impaired mice revealed that activity levels were lower in both CA1 and CA3 in memory-impaired mice 1 month after memory formation, indicating an important role of these areas in early remote memory retrieval. At very remote time points, this reduction persisted only in CA1, suggesting a persistent role of CA1 in memory recall independent of its age and a decreasing relevance of CA3 function for this process over time (three-way ANOVA with memory age, performance and regions as factors, all effects: Ps < 0.01; post hoc comparisons for memory-intact versus memory-impaired: CA1, Ps < 0.01; CA3, 1month: P = 0.003, 6-12 months: P = 0.75 ; Bonferroni-Holm corrected). \**P* < 0.05; \*\**P* < 0.01; \*\*\**P* < 0.001. Bars represent mean +-SEM. **D)** Schematic summary of the most relevant patterns of recruitment of entorhinal, prefrontal and hippocampal subregions at test in memory-intact and memory-impaired mice at recent (1 day or 1 week), early remote (1 month), and very remote (6 and 12 months) time points. Full colors indicate neuronal engagement during the test phase. Lighter colors and grey/no color indicate significantly lower activity or no recruitment, respectively. To facilitate interpretation, only patterns with >5% task-induced *Arc*⁺ expression are presented.

Next, to assess whether LEC and MEC’s functional integrity was required to recall remote memory, we compared activity levels between memory-intact and memory-impaired mice across areas and remote delays (Fig. 2A). The proportion of *Arc+* cells was reduced only in LEC in memory-impaired vs memory-intact mice for early remote memory recall, while it was reduced for both areas at very remote delays (three-way ANOVA, delay × region × performance interaction effect: F(1,14) = 10.55, *P* = 0.006; main performance and delay effects: *Ps* < 0.04, main region effect: *P* = 0.92; post hoc comparisons: memory-impaired vs memory-intact, 1 month: LEC: *P* = 0.04, MEC: *P* > 0.99; 6-12 months: LEC or MEC: *Ps* < 0.01, Bonferroni-Holm corrected; Fig. 2A). This subregional functional segregation was further supported by within-delay analyses across areas reporting a significant region × performance interaction effect at the early remote time-point (1 month groups: two-way-ANOVA: F(1,6) = 10.24, *P* = 0.02, main performance effect: *P* = 0.004, main region effect: *P* = 0.31) but not when the memory was more remote (6-12 months groups: two-way ANOVA: F(1,8) = 0.43, *P* = 0.53, main performance and region effects: *Ps* < 0.005). Of note, baseline *Arc* expression of age-matched controls in LEC or MEC was comparable across delays (two-way ANOVA, delay × area interaction effect: F(4,15) = 0.629; *P* = 0.649, other effects *Ps* > 0.685), hence can unlikely account for the latter differences. Thus, our results indicate a division of roles between the LEC and MEC in their contributions to memory consolidation over time. Although both the LEC and MEC are engaged during object-location recall regardless of memory age, only LEC’s functional integrity is required for recalling memory once they become remote (early remote memories). In contrast, the intact functioning of both the LEC and MEC is ultimately necessary for the recall of memories as they continue to age (very remote memories).

Next, given the well-established role of the PFC in memory consolidation, we aim at distinguishing the specific contributions of its individual subregions within this framework and ultimately compare them with those of the EC subregions.

### Within the PFC, the ACC contributes the most to object-location memory recall. Its recruitment is essential for recalling the oldest memories

Earlier studies have underlined the role of the PFC for the retrieval of early remote fear memories in rodents (i.e., 1-month-old memory^35–37^). However, its role in recalling more remote memories -such as those as old as the memories commonly investigated in human memory research-has been scarcely studied. Moreover, the specific role of its subareas (ACC, PL and IL) in system memory consolidation is not well understood.

First, we evaluated the contribution of the ACC, PL and IL to the successful retrieval of memory over time (memory-intact groups: orange, yellow and brown bars, respectively; Fig. 2B). Statistical analyses of the proportion of task-induced *Arc*+ cells in memory-intact mice showed that the ACC was engaged (or showed a strong tendency thereof) independently of the age of the memory, while the PL and the IL were not significantly recruited in this task (memory-intact groups: comparison to 0: ACC: 1 and 6-12 months: *Ps* < 0.015, 1 day and 1 week: *Ps* < 0.075; PL: *Ps* > 0.281; IL: *Ps* > 0.133; Bonferroni-Holm corrected).

Between PFC areas comparisons further supported this functional segregation (memory-intact groups: two-way ANOVA, area × delay interaction effect: F(6,24) = 5.24, *P* < 0.01; main delay and region effects: *Ps* < 0.017). Indeed, post hoc analyses revealed that the proportion of task-induced *Arc*+ cells were significantly larger in the ACC than in the IL or PL at the remote time-points while it remained similar between areas at the recent ones (ACC vs IL or PL: 1 month- or 6-12 months: *Ps* < 0.039; 1 day or 1 week-: *Ps* > 0.99; Bonferroni-Holm corrected). In addition, within areas analyses showed that ACC activity levels were significantly enhanced for the recall of remote memories compared to that of recent memories, which was not the case for the IL and PL (memory-intact groups: 1 month or 6-12 months-old vs 1 day or 1 week-old: ACC: *Ps <* 0.018; IL and PL: *Ps* > 0.99; Bonferroni-Holm corrected). Of note, here also baseline *Arc* expression in age-matched controls was comparable between PFC regions (two-way ANOVA, area × delay interaction effect: F(8,30) = 0.23, *P* = 0.982, main delay effect *P* = 0.747, main region effect *P* < 0.001), hence is unlikely to account for the differences in brain patterns observed during memory recall overtime.

In summary, these results indicate that, when object-location memory is successfully recalled, the ACC is engaged for recalling memories independently of its age and that this engagement is maximal for the recall of remote memories. In contrast, the contribution of the IL and the PL to this process over time appears negligeable.

Next, we investigated the extent to which PFC areas recruitment was truly necessary to recall remote memories (early and very remote) by comparing brain activity patterns between memory-intact and memory-impaired mice. A comparison of the levels of activity across areas and delays confirmed the existence of a functional segregation within the PFC for the recall of remote memories (three-way ANOVA, performance × delay × area interaction effect: F(2,28) = 21.31; *P* < 0.001; performance and region main effects *Ps* < 0.001; main delay effect *P* = 0.251). Further within area analyses, indicated that ACC activity levels were higher in memory-intact than memory-impaired-groups during very remote but not during early remote memory recall, suggesting a prevalent role of the ACC for the retrieval of the most remote memories (two-way ANOVA, ACC: performance × delay interaction effect: F(1,14) = 15.31, *P* = 0.002, other main effects: *Ps* < 0.02; post hoc comparisons, impaired vs intact: 6-12 month-old: *P* < 0.001, 1 month-old: *P* = 0.37; Bonferroni-Holm corrected). In line with this finding, task-induced ACC *Arc* expression was negligible in mice failing to retrieve the object-location memory only when the oldest memories were recalled (memory-impaired mice: comparisons to 0, 1 month: *P* = 0.046; 6-12 months: *P* = 0.091; Bonferroni-Holm corrected). In contrast, IL or PL activity levels were very low in both memory-impaired and memory-intact mice during early and very remote memory recall, suggesting a lesser involvement of these PFC areas in recalling remote object-location memory (comparison to 0: all *Ps* > 0.133 but memory-impaired: PL: 1 month:*P* = 0.009 and 6-12 month: *P* = 0.047 and IL: 6-12 months:*P* = 0.002; Bonferroni-Holm corrected). Altogether these data reveal that, among the PFC areas, the IL and PL might not play a crucial role in recalling memories over time. In striking contrast, the ACC contributes to recalling memory for the location of objects independently of its age, is maximally engaged when memories become remote, and its functional integrity is ultimately required for very remote memory recall.

Next, we sought to compare directly the patterns of activity in the prefrontal and MTL cortical areas that we identified as relevant and/or necessary for the recall of remote memories. To do so, we compared the proportions of *Arc*+ cells in the ACC, LEC and MEC between memory-impaired and memory-intact mice.

### The LEC is already necessary for recalling early-remote object-location memories, while the MEC and ACC become critical only when memories become more remote

The comparison of the proportions of *Arc+* cells in the ACC, LEC, and MEC between memory-impaired and memory-intact mice during early-remote and very-remote memory recall revealed that patterns of brain activity differed across cortical areas over time (three-way ANOVA, area × performance × delay interaction effect: F(2,28) = 7.71, *P* = 0.002; all other effects: *P* < 0.014). Moreover, post hoc comparisons indicated that, for the ACC and MEC, activity levels differed between intact and impaired groups only for the most remote memories (MEC or ACC: very remote, *Ps* < 0.001; early remote, *Ps* > 0.721; Bonferroni-Holm corrected), indicating that the timing of their involvement in memory recall over time is comparable.

In contrast to the ACC, LEC task-induced *Arc* expression was lower in impaired than in intact groups both at the early-remote time point (*P* = 0.007) and at the very-remote time point (*P* < 0.001; Bonferroni-Holm corrected), suggesting that remote memory recall is first sensitive to LEC dysfunction and becomes sensitive to ACC dysfunction when memories are more remote, as it was the case when comparing the LEC and MEC.

Thus, the LEC might have a predominant role over time in retrieving the location of objects compared to the ACC and MEC, while the latter areas become necessary to this process as the memory becomes increasingly remote. Next, given that the fact that recent studies indicate that hippocampal subfields CA1 and CA3 make distinct contributions to memory consolidation over time, we examined whether such differences are reflected in the present task using the same analytical approach as described above.

### Unlike CA1, CA3’s contribution to the recall of object-location memory is temporally restricted

We first investigated the roles of CA1 and CA3 in successful memory recall by examining brain activity patterns in memory-intact mice (bright and dark red bars, Fig. 2C). A direct between-areas comparison of the proportions of *Arc*+ cells over time showed that CA1 and CA3 were differentially recruited as memories aged, and that CA3 was selectively less engaged than CA1 during the recall of the oldest memories (memory-intact groups: two-way ANOVA, delay × area interaction effect: F(3,12) = 21.9; *P* < 0.001; main delay and area effects: *Ps* < 0.004; post hoc comparisons CA1 vs CA3: recent or early remote groups: *Ps* > 0.882; very remote: *P* = 0.046; Bonferroni-Holm corrected).

Further within-area analyses confirmed this functional segregation by revealing that, unlike CA3, CA1 was recruited to a similar extent during memory recall regardless of memory age (memory-intact groups: one-way ANOVAs, CA1: F(3,12) = 0.48; *P* = 0.700; CA3: F(3,12) = 21.3; *P* < 0.001; Tukey post hoc tests: CA3: very remote vs recent or early remote: *Ps* < 0.001; Fig. 2C).

In addition, both CA1 and CA3 were significantly activated during the recall of recent and early remote memories, whereas only CA1 remained engaged when memories were more remote (memory-intact groups: comparisons against 0: CA1 or CA3, recent and early remote: *Ps* < 0.01; very remote: CA1: *P* = 0.002; CA3: *P* = 0.132; Bonferroni-Holm corrected; Fig. 2C). Importantly, baseline *Arc* expression in age-matched controls was comparable between CA1 and CA3 and across delays, indicating that constitutive differences between areas or across delays are unlikely to account for the differences in activity patterns observed during memory recall (two-way ANOVA, area × delay interaction effect: F(4,15) = 0.339; *P* = 0.848; main delay effect: *P* = 0.689, main region effect *P* = 0.021). Altogether, these results indicate that CA3 engagement during object-location recall is limited to recent and early remote memories, whereas the contribution of CA1 is independent of memory age.

Next, we investigated whether CA1 or CA3 were necessary for memory recall by comparing activity levels between memory-intact and memory-impaired mice over time. These analyses revealed differences in the roles of CA1 and CA3 in remote memory recall as memories aged (three-way ANOVA, performance × area × delay interaction effect: F(1,14) = 21.38; *P* < 0.001; other effects *Ps* < 0.010 but performance × delay interaction effect: *P* = 0.29).

Further within-delay comparisons between intact and impaired memory groups showed that CA1 and CA3 differed functionally specifically when memories were very remote (two-way ANOVAs, early remote: area × performance interaction effect: F(1,6) = 0.91, *P* = 0.378; very remote: area × performance interaction effect: F(1,8) = 26.15, *P* < 0.001; other effects *Ps* < 0.010; post hoc comparisons, intact vs impaired: CA1: *P* < 0.001; CA3: *P* = 0.75; Bonferroni-Holm corrected).

This result was further supported by within-area analyses showing that the proportion of *Arc+* cells in CA1 was reduced in memory-impaired compared with memory-intact mice during both early and very remote memory recall to a similar extent (two-way ANOVA, delay × performance interaction effect: F(1,14) = 4.129; *P* = 0.061; main performance effect: *P* < 0.001; main delay effect: *P* = 0.3). In contrast, such a reduction was observed only during early remote memory recall in CA3 (two-way ANOVA, delay × performance interaction effect: F(1,14) = 17.47; *P* < 0.001; main performance and delay effects: *Ps* < 0.010; post hoc comparisons intact vs impaired: early remote: *P* = 0.003, very remote: *P* = 0.75, Bonferroni-Holm corrected).

Altogether, our results indicate that both CA1 and CA3 contribute to, and are necessary for the recall of early remote object-location memories. In addition, these findings show that the functional integrity of CA1 is also required for retrieving more remote memories, whereas this is no longer the case for CA3 at this time point. Thus, CA1 appears to play a predominant and persistent role in the recall of remote object-location memory, whereas CA3’s contribution appears to be temporally limited.

Overall, our findings reveal a time-dependent, circuit-specific reorganization of the neural circuits supporting object-location memory recall and identify the EC as an important contributor to memory consolidation alongside the PFC. In the hippocampus, both CA1 and CA3 contribute to the retrieval of early remote memories, but only CA1 remains necessary as memories become more remote. In parallel, within the entorhinal cortex, both LEC and MEC are engaged during recall across delays; however, LEC’s functional integrity is already required to recall early remote memory, whereas MEC becomes necessary only when memories are very remote. At the PFC level, the ACC is selectively engaged during memory recall regardless of memory age, with stronger recruitment for remote memories. Notably, MEC and ACC exhibit similar temporal dynamics, with both regions showing increased relevance as memories become very remote.

## Discussion

The present study provides new insights into the systems-level organization of long-term memory by revealing how distinct cortico-hippocampal circuits are differentially engaged during the recall of early and very remote memories. Using a novel version of the object-location task, we demonstrate that memory traces are not diffusely redistributed across the cortex over time but instead follow an organized and temporally structured reorganization involving specific subregions of the entorhinal cortex (EC), prefrontal cortex (PFC), and hippocampus (HIP). By exploiting the natural decay of memory, we causally dissociated the contribution of these regions to remote memory recall and uncovered a temporal shift from a LEC-CA1/CA3 circuit supporting early remote memory recall to a distinct more distributed LEC/MEC-ACC-CA1 network supporting more remote memories. This reconfiguration features a time-dependent functional segregation within the hippocampus, the EC and the PFC, suggesting that hippocampal involvement in memory recall evolves in parallel with cortical reorganization. Together, these findings refine current systems consolidation theories by demonstrating that memory consolidation is both circuit-specific and temporally ordered, and by positioning the entorhinal cortex as an important dynamic component of long-term memory retrieval, together with the PFC.

Disentangling the contribution of entorhinal cortex (EC) subregions to memory recall has so far relied largely on rodent studies, in part because the homologous functional subdivisions of the human EC have been identified with MRI only relatively recently ^38,39^. In both species, prior work mainly focused on the recall of recent memories and/or the role of EC subareas in spatial or non-spatial information processing ^11,16,18,20,21,40^, rather than on their involvement in memory consolidation over time. By contrast, only few studies have directly examined the role of EC subregions in early remote memory consolidation, i.e. up to one month following memory formation ^23,41^. Moreover, even fewer have adopted causal approaches ^24^ or time-windows of investigation comparable to those used in human studies ^23,24^, limiting translational inference between species. As a result, whether EC subareas contribute equivalently to remote memory recall over time - or whether they are truly necessary for this process - has remained poorly understood. Our results show that both LEC and MEC are robustly recruited during remote memory recall, assessed for object-location associations. These results are in line with recent findings obtained using contextual fear-conditioning paradigms ^23,24^, which, however, did only investigate the level of engagement of these areas but not whether the LEC and MEC’s functional integrity were necessary for this process. Here, by combining neural activity mapping with a causal approach across extended time scales, we extend prior knowledge on the subject by revealing a temporal dissociation in the functional contributions of the EC subregions. Indeed, while both LEC and MEC exhibit maximal engagement during remote recall, only the functional integrity of the LEC is required for the recall of early remote memories (1 month-old), as evidenced by a pronounced reduction in LEC activity in memory-impaired mice as compared to memory-intact mice. In contrast, recall of more remote memories (6-12 months old) depends on the integrity of both MEC and LEC areas, as shown by activity levels reduced in both subregions in animals exhibiting deficits. Together, these findings identify a time-dependent reorganization within the EC, highlighting distinct and evolving roles of its subregions in supporting remote memory recall, with a primary role of the LEC in recalling early remote memories and both the LEC and MEC contributing to the recall of older memories.

In a similar line, our data reveal a robust and evolving functional segregation across PFC subregions in their contribution to memory consolidation over time. Notably, the ACC was selectively recruited during remote memory recall, whereas the IL and PL showed only minimal involvement, both in temporal engagement and response magnitude. This observation is noteworthy given prior evidence implicating the PL and IL in the strengthening of fear memory in the consolidation and extinction of recent emotional memories ^42–44^, as well as anatomical data indicating that hippocampal projections to the PFC preferentially target the PL^42,45,46^. Despite these findings, activity in the IL and PL of mice expressing intact memory traces did not reach statistical significance here, i.e. when the recall of remote object-location memories was investigated. Collectively, these results might suggest that IL and PL contributions to retrieving more neutral memories (object-location) are comparatively limited relative to their established roles in emotional memory processing. In striking contrast, we observed robust ACC engagement, during early remote memory recall (1-month old), in agreement with the systems-consolidation literature. At this time-point, the ACC is thought to coordinate activity across distributed neuronal ensembles and cortical regions in which distinct components of a memory are stored ^3,47,48^. We extend this finding by demonstrating ACC involvement during the recall of much older memories (6-12 months old), a temporal window comparable to that typically examined in humans but largely unexplored in rodent models, thereby strengthening translational inferences across species. Importantly, our causal analyses further reveal that although the ACC is already strongly engaged during early remote object-location memory recall, its activity becomes functionally necessary only at later stages. Specifically, reduced ACC activity was observed exclusively in memory-impaired mice during recall of the oldest memories, whereas ACC engagement persists in mice with memory deficits at the early remote time point. This apparent deviation from classical systems-consolidation models may, at least in part, reflect differences in memory domain. Much of the systems-consolidation literature is based on fear-conditioning paradigms, in which consolidation is known to occur rapidly ^3^. By contrast, the consolidation of more neutral memories, such as object-location associations, is thought to unfold more slowly, potentially due to their lower emotional salience and reduced immediate relevance for survival. Accordingly, dependence on ACC function for the retrieval of neutral associations may emerge at later time points than for emotional or aversive memories, as observed here. Together, our findings reveal that PFC subregions make domain-specific and temporally distinct dynamic contributions to long-term memory retrieval and highlight the role of the ACC in the recall of memories comparatively as old as those studied in remote memory research in humans.

Notably, the temporal profile of ACC involvement in remote memory recall closely parallels that observed for the MEC, suggesting that these regions may participate in a shared functional network that becomes particularly relevant for the retrieval of very remote memories. This possibility is supported by anatomical evidence revealing robust polysynaptic connectivity between the MEC and ACC via intermediary structures including the hippocampus, retrosplenial cortex, and nucleus reuniens, alongside relatively sparse direct projections ^49,50^.

Furthermore, our data suggest that the putative LEC/MEC-ACC network, necessary for very remote memory recall, might selectively include the CA1 subfield of the hippocampus. Specifically, activity levels were reduced in CA1, but not CA3, in memory-impaired mice during retrieval of very remote object-location memories. This interpretation is consistent with previous IEG imaging and optogenetic fear-conditioning studies demonstrating persistent CA1 recruitment for memory recall, even at very remote time points, in which CA1 activity together with MTL cortical activity was proposed to support the retrieval of generalized or gist-like memory representations ^24^. However, these latter studies did not causally test the contribution of EC subregions and did not examine PFC regions. In addition, these results are in line with the electrophysiological reports in rodents of increased ACC-CA1 theta coherence during (early) remote recall ^51^ and findings in humans according to which functional disruption of CA1 in transient global amnesia patients impairs even the memory for 30 to 40 years old events ^52^. Importantly, in contrast to CA1, we observed that although CA3 contributes to retrieval at early remote time points, it is no longer required for recalling older memories. This finding is consistent with optogenetic fear-conditioning data indicating that CA3 is selectively required for retrieval of precise memory representations up to approximately one month after encoding ^23,24^. More broadly, the sustained involvement of CA1 coupled with the temporally restricted contribution of CA3 - here extended to more neutral object-location memories - suggests that this division of labor might reflect a fundamental principle for systems-level memory consolidation. Together, these latter findings show that CA3 is not a core component of the subnetwork, including the LEC/MEC, ACC and CA1, that supports retrieval of the most remote memories. Importantly, while our data reveal a time-limited contribution of CA3 and a sustained involvement of CA1 for the recall of remote memories, the causal roles of CA1 and CA3 during *recent* memory recall could not be directly assessed in the present study as mice did not exhibit memory deficits at these earlier time points. However, this aspect falls outside the scope of the present study, which focuses on remote memory recall, and has been extensively addressed by previous work demonstrating critical contributions of both CA1 and CA3 to recent memory retrieval in a causal manner^53–58^.

Taken together, our results suggest a progressive shift from a network primarily involving the LEC and hippocampal CA1/CA3 subfields toward a distinct and more distributed network encompassing the LEC/MEC, ACC, and CA1 during retrieval of object-location memories as they age. This transition is consistent with known functional specializations within the medial temporal lobe as the medial entorhinal cortex (MEC) is predominantly associated with spatial processing, whereas the lateral entorhinal cortex (LEC) is more strongly linked to object-related and contextual information ^11,16,17,40^, and both CA1 and CA3 have been implicated in object-location memory processing ^9,11^. Within this framework, the reliance of early remote memory recall on the functional integrity of LEC, CA1, and CA3 might reflect task-dependent memory demands. Specifically, retrieval of object-location associations may preferentially engage circuits specialized for representing these features as long as the memory retains sufficient representational precision. As memory representations become more remote and lose details^24,59,60^, one can speculate that residual mnemonic information may increasingly overlap with representational formats supported by other cortical circuits. Consequently, retrieval may become progressively supported by regions that are not primarily specialized for the original informational content, but that can nevertheless process remaining shared or partially preserved features of the representation. Consistent with this interpretation, ACC and MEC activity levels during early remote recall were considerable and comparable between memory-impaired and memory-intact mice, suggesting that recruitment of these regions at this stage may not yet reflect functionally effective processing of object-location information. Instead, these regions - whose canonical functions are not primarily object-location specific - might become more functionally relevant as memory representations lose precision over time, thereby enabling engagement of a broader, more distributed network. Testing this hypothesis is beyond the scope of the present study and will require further investigations

Altogether, by examining memory across time windows that enhance translational relevance to humans, we reveal a temporal shift from a LEC-CA1/CA3 circuit supporting early remote object-location memory recall to a distinct, more distributed LEC/MEC-ACC-CA1 network critical for older memories. These findings refine current systems consolidation theories by showing that network reorganization is not only time-dependent but might also be circuit-specific, likely shaped by the representational content of the memory. Moreover, our results highlight the entorhinal cortex as a dynamic and central component of long-term memory retrieval, alongside the PFC.

## Material and Methods

### Ethics statement

All experimental procedures were approved by the State of Saxony-Anhalt ethics committee under license 4252-2 1555 LIN and carried out in accordance with the European Communities Council Directive of September 22^nd^, 2010 (2010/63/EU).

### Animals

Male C56BL/6 mice (10 – 70 weeks old; n = 55 bred at the Leibniz Institute for Neurobiology (Magdeburg, Germany) were used. Animals were group housed and allowed to acclimate to the housing facility for 1 week, and were single housed at least 48 h prior to the start of behavioral procedures. Mice were maintained under a reversed 12 h light/dark cycle (lights off at 7 a.m.; lights on at 7 p.m.) to enable testing during their active phase. Food and water were available ad libitum.

Sample sizes (n = 4-8 per group) were determined based on previous studies^9,23,24,30^. Mice were randomly assigned to experimental groups prior to the start of the experiments. Experimental groups were defined by the delay between memory formation and testing (1 day, n = 4; 1 week, n = 4; 1 month, n = 8; 6 months, n = 6; 12 months, n = 4). Age-matched control mice (n = 20) that were place in the same experimental room but did not undergo testing were included in the study (see Habituation and Testing schedule). Experimental animals that did not reach behavioral criteria (n = 9) were excluded from further analyses (see Behavioral analysis).

### Behavioral paradigm

**Apparatus and stimuli.** Behavioral testing was conducted in a 32 × 32 × 41 cm open field, placed in a dimly lit room. Extra-maze cues, as well as cues on the outside wall of the open field, served as spatial references. A video camera (Sony, HDR/CX500E) recorded the animals’ behavior for off-line analysis. Four copies of 2 different metallic objects were used for testing so that objects used during the test phase were duplicates of those during the study phases.

**Habitation and testing**. Habituation procedure occurred on 4 consecutive days following a procedure similar to Beer et al., 2014. In brief, animals were habituated to the empty open field for 20 min during days 1 and 2. To encourage mice to explore all areas of the box (divided in 9 quadrants) and minimize the development of a spatial bias, 1 chocolate sprinkle was placed in the center of each quadrant. The absence of displacement of sprinkles and/or the presence of droppings in each quadrant at the end of each 20-min session were assessed and revealed that mice had explored each quadrant during each session. On days 3 and 4, animals were habituated to the presence of objects, which were not used on the testing day in conditions that mimicked those of the testing day (one15-min trial followed by a 6-min trial each day). Across habituation sessions, object positions varied to ensure full exploration of the arena. After each trial the open field and stimuli were cleaned using water and a solution containing 10% ethanol.

The testing procedure was adapted from previous studies on spontaneous object recognition tasks ^9,11,32,62^. In brief, to enable the study of very remote memories the duration of the study phase was extended to 15 min as performed in Melani et al. 2017^63^. To accommodate the detection of *Arc* mRNA the test phase was shortened to 6 min^9,11,27,64^. During the study phase, mice explored two identical objects placed in the open field and were returned to their home cage. After a retention interval (1 day, 1 week, 1 month, 6 months, or 12 months), mice were reintroduced to the arena for a 6 min test phase. During testing, duplicates of the study objects were placed in the arena: one object remained in its original location (the ‘stationary’ object), whereas the other was displaced (see Fig. 1A). The position of the objects during the study and test was counterbalanced across animals. After each trial the open field and stimuli were cleaned using water and a solution containing 10% ethanol.

Age-matched control mice were brought to the experimental room but did not undergo habituation or testing procedures (i.e. remained in their home-cage). These mice were used to evaluate *Arc* mRNA expression due to parameters other than task demands (baseline *Arc* expression) and calculate the task-induced *Arc* _expression_11,25,29,31,64,65

### Behavioral analysis

Based on animal’s natural preference for novelty^32^, a successful memory for a given spatial location is observed when the “displaced object” is explored more than the “stationary object”. The exploration time for an object was defined as the time spent in exploring an object, i.e., nose at a distance <2cm to the object as originally described in Ennaceur and Delacour, 1998^32^. Behavioral performance was scored off line by 2 independent experimenters blind to experimental conditions and averaged. The discrimination index (DI) during the test phase was calculated as follows: DI = [(Exploration Time for the displaced object) – (Exploration Time for the stationary object)]/ (Total Exploration Time). Animals that did not explore each object for at least 10 sec during the study phase were excluded (n = 9)^66^. It was ensured that exploration of the objects during the study phase were comparable across groups.

**Classification of memory-intact and memory-impaired mice.** To classify animals based on behavioral performance, unsupervised K mean clustering was performed on discrimination indices (DI) obtained during the test phase. Clustering was performed on individual DI values without prior labeling. The algorithm was initialized using random initialization (with multiple random starts, nstart = 25), and cluster assignment was based on minimization of within-cluster variance which yielded a threshold of 0.09. Resulting clusters were interpreted post hoc based on their mean DIs, with the cluster exhibiting. higher values classified as ‘memory-intact’ (mean DI = 0.33 ± 0.12; MEAN±SEM) and the lower DI as ‘memory-impaired’ mice (mean DI = −0.11 ± 0.19; MEAN±SEM).

**Imaging activity in the hippocampus, entorhinal, frontal cortical areas.** To assess the neural correlates of memory retrieval across recent, and remote delays, brains of experimental and control animals were processed for the detection of the pre-mRNA of the Immediate-Early-Gene (IEG) *Arc* by *in-situ* hybridization to evaluate the percentage of cells recruited at test for each targeted brain area. This technique yields cellular resolution, allowing precise evaluation of recruitment levels in adjacent brain areas such as CA1, CA3, LEC, and MEC^23,25^. The detection of the expression of the IEG *Arc* was preferred over that of other IEGs such as *Fos* and *zif 268* because it has strong ties to synaptic plasticity, reflects better memory demands and not stress levels ^20,27,29^ and has been heavily used for mapping memory-like activity in the hippocampus and the parahippocampal areas for the past decades. Following the standard protocol described in Beer et al., 2013, *Arc* pre-mRNA was detected so that only *Arc* intranuclear signal was observable ^9,11,25,61,67^.

In short, mice were decapitated upon the recognition phase of the object-in-place task. Brains were removed, flash frozen in isopentane, and stored at −80°C until sectioning. Brains were sectioned with a cryostat (Leica CM 3050 S; 8-μm-thick coronal sections), mounted on Polylysine slides (Thermo Scientific), and stored at −80°C until *in-situ* hybridization. *Arc* pre-mRNA probes were synthesized using the digoxigenin-labeled UTP kit (Roche Diagnostics). Slides were fixed with 4% buffered paraformaldehyde and rinsed with 0.1 M PBS. Slides were treated with an acetic anhydride/triethanolamine/hydrochloric acid mix, rinsed, and briefly soaked with a prehybridization buffer. The tissue was hybridized with the digoxigenin-labeled *Arc* probe overnight at +65°C. Following hybridization, slides were rinsed with buffer solutions and treated with an antidigoxigenin -horseradish peroxidase (HRP) conjugate (Roche Molecular Biochemicals) and a cyanin-3 substrate kit (Cy3, TSA-Plus system, Perkin Elmer). Nuclei were counterstained with 4’, 6’-diamidino-2-phenylindole (DAPI; Vector Laboratories). To detect *Arc* signal in all target areas, three slides per animal were processed and analyzed. Slides contained four nonconsecutive brain section (distant ca 50 μm). One slide contained the prefrontal areas (AP 1.7 mm), the second hippocampal areas and LEC (AP −3.0 mm) and the third the MEC (AP −4.5 mm) (Fig. 1B;^68^). In agreement with standard IEG imaging protocols, images from three nonadjacent sections for each area of interest were acquired that covered approximately 200 μm. The number of activated neurons was evaluated on approximately 50-90 neurons per image. Images were captured with a Keyence Fluorescence microscope (BZ-X710; Japan). Images were taken with a 40× objective (0.7-μm -thick z-stacks). Exposure time and light intensity were kept similar for image acquisition across all slides. As first described in the seminal work of Guzowski and colleagues ^25^, contrasts were set to optimize the appearance of intranuclear foci ^9,11,61,67^. To account for stereological considerations, neurons were counted on 8-μm-thick sections that contained 1 layer of cells, and only cells containing whole nuclei were included in the analysis ^69^. The quantification of *Arc* expression was performed in the median 60% of the stack because this method minimizes the likelihood of taking into consideration partial nuclei and decreases the occurrence of false negatives. This method is comparable to an optical dissector technique that reduces sampling errors linked to the inclusion of partial cells into the counts and stereological concerns because variations in cell volumes no longer affect sampling frequencies ^69^. Also, as performed in a standard manner in *Arc* imaging studies, counting was performed on cells (>5 μm) thought to be pyramidal neurons or interneurons because small non-neuronal cells such as astrocytes or inhibitory neurons do not express *Arc* following behavioral test^38^. The designation “intranuclear-foci-positive neurons” (*Arc*-positive neurons) was given when the DAPI-labeled nucleus of the presumptive neurons showed 1 or 2 characteristic intense intranuclear areas of fluorescence. DAPI-labeled nuclei that did not contain fluorescent intranuclear foci were counted as “negative” (*Arc*-negative neurons) ^25^. Percentage of *Arc-*positive neurons was calculated as follows: *Arc*-positive neurons / (*Arc*-positive neurons + *Arc*-negative neurons) × 100. To isolate *Arc* expression induced by memory retrieval, data were normalized by subtracting baseline *Arc* levels measured in age-matched control groups.

**Statistical Analysis.** K mean clustering was used to allocate mice into memory-intact and memory impaired-groups (MATLAB). All statistical analyses were performed using RStudio (version 2025.05.1). For behavioral data and task-induced *Arc* expression, differences from 0 were assessed using two-tailed one-sample t-tests. For behavioral analyses, group comparisons between memory-intact and memory-impaired mice or comparisons across delays were performed using one-way or two-way ANOVAs (analysis of variance) and post hoc two-tailed unpaired t-tests with Bonferroni-Holm correction for multiple testing. For raw or task-induced *Arc* expression analyses, one-, two- or three-way ANOVAs were applied with performance (memory-intact vs memory-impaired) and delay (1 day, 1 week, 1 month, 6-12 months) as between-subject factors, and region (CA1, CA3, ACC, PL, IL, LEC, MEC) as a within-subject factors were appropriate. Multiple comparisons were corrected using Bonferroni-Holm or Tukey tests. All tests were two-tailed, and statistical significance was defined as P < 0.05.

## Acknowledgments

Funded by the German Federal Ministry of Education and Research (BMBF) under grant number 01EE2305E. Funded by the Deutsche Forschungsgemeinschaft (DFG, German Research Foundation) - CRC1436

## References

1. Alvarez, P. & Squire, L. R. Memory consolidation and the medial temporal lobe: A simple network model. Proc. Natl. Acad. Sci. U. S. A. 91, 7041–7045 (1994).

2. Nadel, L. & Moscovitch, M. Memory consolidation, retrograde amnesia and the hippocampal complex. Curr. Opin. Neurobiol. 7, 217–227 (1997).

3. Frankland, P. W. & Bontempi, B. The organization of recent and remote memories. Nat. Rev. Neurosci. 6, 119–130 (2005).

4. Atucha, E. et al. Noradrenergic activation of the basolateral amygdala maintains hippocampus-dependent accuracy of remote memory. Proc. Natl. Acad. Sci. U. S. A. 114, 9176–9181 (2017).

5. Yonelinas, A. P., Ranganath, C., Ekstrom, A. D. & Wiltgen, B. J. A contextual binding theory of episodic memory: systems consolidation reconsidered. Nature Reviews Neuroscience vol. 20 364–375.

6. Kitamura, T. et al. Engrams and circuits crucial for systems consolidation of a memory. Science (1979). 356, 73–78 (2017).

7. Moscovitch, M. & Gilboa, A. Systems consolidation, transformation and reorganization: Multiple Trace Theory, Trace Transformation Theory and their Competitors in Michael J. Kahana, and Anthony D. Wagner (eds). The Oxford Handbook of Human Memory, Two Volume Pack: Foundations and Applications, Oxford Library of Psychology (2024)

8. van Strien, N. M., Cappaert, N. L. M. & Witter, M. P. The anatomy of memory: an interactive overview of the parahippocampal–hippocampal network. Nat. Rev. Neurosci. 10, 272–282 (2009).

9. Beer, Z., Chwiesko, C. & Sauvage, M. M. Processing of spatial and non-spatial information reveals functional homogeneity along the dorso-ventral axis of CA3, but not CA1. Neurobiol. Learn. Mem. 111, 56–64 (2014).

10. Leutgeb, S., Leutgeb, J. K., Treves, A., Moser, M. B. & Moser, E. I. Distinct ensemble codes in hippocampal areas CA3 and CA1. Science (1979). 305, 1295–1298 (2004).

11. Beer, Z., Chwiesko, C., Kitsukawa, T. & Sauvage, M. M. Spatial and stimulus-type tuning in the LEC, MEC, POR, PrC, CA1, and CA3 during spontaneous item recognition memory. Hippocampus 23, 1425–1438 (2013).

12. Sauvage, M. M. Neural Substrates of Recollection and Familiarity: Further Bridging Human and Animal Recognition Memory Using Translational Paradigms. Psychology of Memory (2012).

13. Heidbreder, C. A. & Groenewegen, H. J. The medial prefrontal cortex in the rat: evidence for a dorso-ventral distinction based upon functional and anatomical characteristics. Neurosci. Biobehav. Rev. 27, 555–579 (2003).

14. Laubach, M., Amarante, L. M., Swanson, K. & White, S. R. What, If Anything, Is Rodent Prefrontal Cortex? eNeuro 5, ENEURO.0315-18.2018 (2018).

15. Dalley, J. W., Cardinal, R. N. & Robbins, T. W. Prefrontal executive and cognitive functions in rodents: neural and neurochemical substrates. Neurosci. Biobehav. Rev. 28, 771–784 (2004).

16. Fyhn, M., Molden, S., Witter, M. P., Moser, E. I. & Moser, M.-B. Spatial Representation in the Entorhinal Cortex. Science (1979). 305, 1258–1264 (2004).

17. Save, E. & Sargolini, F. Disentangling the role of the MEC and LEC in the processing of spatial and non-spatial information: Contribution of lesion studies. Frontiers in Systems Neuroscience vol. 11.

18. Deshmukh, S. S. & Knierim, J. J. Representation of non-spatial and spatial information in the lateral entorhinal cortex. Front. Behav. Neurosci. 5, (2011).

19. Ku, S.-P. et al. Regional specific evidence for memory-load dependent activity in the dorsal subiculum and the lateral entorhinal cortex. Front. Syst. Neurosci. 11, (2017).

20. Atucha, E., Karew, A., Kitsukawa, T. & Sauvage, M. M. Recognition memory: Cellular evidence of a massive contribution of the LEC to familiarity and a lack of involvement of the hippocampal subfields CA1 and CA3. Hippocampus 27, 1083–1092 (2017).

21. Mahnke, L., Atucha, E., Pina-Fernàndez, E., Kitsukawa, T. & Sauvage, M. M. Lesion of the hippocampus selectively enhances LEC’s activity during recognition memory based on familiarity. Sci. Rep. 11, (2021).

22. Sauvage, M. M., Beer, Z., Ekovich, M., Ho, L. & Eichenbaum, H. The caudal medial entorhinal cortex: A selective role in recollection-based recognition memory. Journal of Neuroscience 30, 15695–15699 (2010).

23. Lux, V., Atucha, E., Kitsukawa, T. & Sauvage, M. M. Imaging a memory trace over half a life-time in the medial temporal lobe reveals a time-limited role of CA3 neurons in retrieval. Elife 5, (2016).

24. Atucha, E., Ku, S.-P., Lippert, M. T. & Sauvage, M. M. Recalling gist memory depends on CA1 hippocampal neurons for lifetime retention and CA3 neurons for memory precision. Cell Rep. 42, 113317 (2023).

25. Guzowski, J. F., McNaughton, B. L., Barnes, C. A. & Worley, P. F. Environment-specific expression of the immediate-early gene Arc in hippocampal neuronal ensembles. Nat. Neurosci. 2, 1120–1124 (1999).

26. Guzowski, J. F., Setlow, B., Wagner, E. K. & McGaugh, J. L. Experience-dependent gene expression in the rat hippocampus after spatial learning: A comparison of the immediate-early genes Arc, c-fos, and zif268. Journal of Neuroscience 21, 5089–5098 (2001).

27. Kubik, S., Miyashita, T. & Guzowski, J. F. Using immediate-early genes to map hippocampal subregional functions. Learning and Memory vol. 14 758–770 (2007).

28. Lux, V., Masseck, O. A., Herlitze, S. & Sauvage, M. M. Optogenetic destabilization of the memory trace in CA1: Insights into reconsolidation and retrieval processes. Cerebral Cortex 27, 841–851 (2017).

29. Nakamura, N. H., Flasbeck, V., Maingret, N., Kitsukawa, T. & Sauvage, M. M. Proximodistal Segregation of Nonspatial Information in CA3: Preferential Recruitment of a Proximal CA3-Distal CA1 Network in Nonspatial Recognition Memory. Journal of Neuroscience 33, 11506–11514 (2013).

30. Sauvage, M. M., Nakamura, N. H. & Beer, Z. Mapping memory function in the medial temporal lobe with the immediate-early gene Arc. Behavioural Brain Research 254, 22–33 (2013).

31. Flasbeck, V., Atucha, E., Nakamura, N. H., Yoshida, M. & Sauvage, M. M. Spatial information is preferentially processed by the distal part of CA3: Implication for memory retrieval. Behavioural Brain Research 354, 31–38 (2018).

32. Ennaceur, A. & Delacour, J. A new one-trial test for neurobiological studies of memory in rats. 1: Behavioral data. Behavioural Brain Research 31, 47–59 (1988).

33. Jain, A. K. Data clustering: 50 years beyond K-means. Pattern Recognit. Lett. 31, 651–666 (2010).

34. Lindenberger, U. Human cognitive aging: *Corriger la fortune?* Science (1979). 346, 572–578 (2014).

35. Frankland, P. W., Bontempi, B., Talton, L. E., Kaczmarek, L. & Silva, A. J. The involvement of the anterior cingulate cortex in remote contextual fear memory. Science 304, 881–883 (2004).

36. Miller, C. A., et al. Cortical DNA methylation maintains remote memory. Nature Publishing Group 13, 664–666 (2010).

37. Vetere, G. et al. Reactivating fear memory under propranolol resets pre-trauma levels of dendritic spines in basolateral amygdala but not dorsal hippocampus neurons. Front. Behav. Neurosci. 7, (2013).

38. Maass, A., Berron, D., Libby, L. A., Ranganath, C. & Düzel, E. Functional subregions of the human entorhinal cortex. Elife 4, (2015).

39. Navarro Schröder, T., Haak, K. V, Zaragoza Jimenez, N. I., Beckmann, C. F. & Doeller, C. F. Functional topography of the human entorhinal cortex. Elife 4, (2015).

40. Knierim, J. J., Neunuebel, J. P. & Deshmukh, S. S. Functional correlates of the lateral and medial entorhinal cortex: objects, path integration and local–global reference frames. Philosophical Transactions of the Royal Society B: Biological Sciences 369, 20130369 (2014).

41. Wheeler, A. L. et al. Identification of a Functional Connectome for Long-Term Fear Memory in Mice. PLoS Comput. Biol. 9, (2013).

42. Giustino, T. F. & Maren, S. The Role of the Medial Prefrontal Cortex in the Conditioning and Extinction of Fear. Front. Behav. Neurosci. 9, (2015).

43. Ye, X., Kapeller-Libermann, D., Travaglia, A., Inda, M. C. & Alberini, C. M. Direct dorsal hippocampal–prelimbic cortex connections strengthen fear memories. Nat. Neurosci. 20, 52–61 (2017).

44. Barker, G. R. I. & Warburton, E. C. NMDA Receptor Plasticity in the Perirhinal and Prefrontal Cortices Is Crucial for the Acquisition of Long-Term Object-in-Place Associative Memory. The Journal of Neuroscience 28, 2837–2844 (2008).

45. Jay, T. M., Glowinski, J. & Thierry, A.-M. Selectivity of the hippocampal projection to the prelimbic area of the prefrontal cortex in the rat. Brain Res. 505, 337–340 (1989).

46. Jay, T. M. & Witter, M. P. Distribution of hippocampal CA1 and subicular efferents in the prefrontal cortex of the rat studied by means of anterograde transport of *Phaseolus vulgaris* -leucoagglutinin. Journal of Comparative Neurology 313, 574–586 (1991).

47. Teixeira, C. M., Pomedli, S. R., Maei, H. R., Kee, N. & Frankland, P. W. Involvement of the anterior cingulate cortex in the expression of remote spatial memory. Journal of Neuroscience 26, 7555–7564 (2006).

48. Wirt, R. & Hyman, J. Integrating Spatial Working Memory and Remote Memory: Interactions between the Medial Prefrontal Cortex and Hippocampus. Brain Sci. 7, 43 (2017).

49. Agster, K. L. & Burwell, R. D. Cortical efferents of the perirhinal, postrhinal, and entorhinal cortices of the rat. Hippocampus 19, 1159–1186 (2009).

50. Varela, C., Kumar, S., Yang, J. Y. & Wilson, M. A. Anatomical substrates for direct interactions between hippocampus, medial prefrontal cortex, and the thalamic nucleus reuniens. Brain Struct. Funct. 219, 911–929 (2014).

51. Wirt, R. A. & Hyman, J. M. ACC Theta Improves Hippocampal Contextual Processing during Remote Recall. Cell Rep. 27, 2313–2327.e4 (2019).

52. Bartsch, T., Döhring, J., Rohr, A., Jansen, O. & Deuschl, G. CA1 neurons in the human hippocampus are critical for autobiographical memory, mental time travel, and autonoetic consciousness. Proc. Natl. Acad. Sci. U. S. A. 108, 17562–17567 (2011).

53. Nakazawa, K. et al. Hippocampal CA3 NMDA receptors are crucial for memory acquisition of one-time experience. Neuron 38, 305–315 (2003).

54. Nakashiba, T., Young, J. Z., McHugh, T. J., Buhl, D. L. & Tonegawa, S. Transgenic inhibition of synaptic transmission reveals role of CA3 output in hippocampal learning. Science 319, 1260–1264 (2008).

55. Dimsdale-Zucker, H. R., Ritchey, M., Ekstrom, A. D., Yonelinas, A. P. & Ranganath, C. CA1 and CA3 differentially support spontaneous retrieval of episodic contexts within human hippocampal subfields. Nat. Commun. 9, (2018).

56. Hunsaker, M. R. & Kesner, R. P. Dissociations across the dorsal–ventral axis of CA3 and CA1 for encoding and retrieval of contextual and auditory-cued fear. Neurobiol. Learn. Mem. 89, 61–69 (2008).

57. Hunsaker, M. R., Thorup, J. A., Welch, T. & Kesner, R. P. The role of CA3 and CA1 in the acquisition of an object-trace-place paired-associate task. Behavioral Neuroscience 120, 1252–1256 (2006).

58. Rolls, E. T. The storage and recall of memories in the hippocampo-cortical system. Cell Tissue Res. 373, 577–604 (2018).

59. Hardt, O., Nader, K. & Nadel, L. Decay happens: The role of active forgetting in memory. Trends in Cognitive Sciences vol. 17 111–120 (2013).

60. Wiltgen, B. J. & Silva, A. J. Memory for context becomes less specific with time. Learning and Memory 14, 313–317 (2007).

61. Vazdarjanova, A. & Guzowski, J. F. Differences in hippocampal neuronal population responses to modifications of an environmental context: Evidence for distinct, yet complementary, functions of CA3 and CA1 ensembles. Journal of Neuroscience 24, 6489–6496 (2004).

62. Aggleton, J. P. Multiple anatomical systems embedded within the primate medial temporal lobe: Implications for hippocampal function. Neurosci. Biobehav. Rev. 36, 1579–1596 (2012).

63. Melani, R., Chelini, G., Cenni, M. C. & Berardi, N. Enriched environment effects on remote object recognition memory. Neuroscience 352, 296–305 (2017).

64. Sauvage, M., Kitsukawa, T. & Atucha, E. Single-cell memory trace imaging with immediate-early genes. J. Neurosci. Methods 326, (2019).

65. Vazdarjanova, A. et al. Spatial exploration induces ARC, a plasticity-related immediate-early gene, only in calcium/calmodulin-dependent protein kinase II-positive principal excitatory and inhibitory neurons of the rat forebrain. Journal of Comparative Neurology 498, 317–329 (2006).

66. Okuda, S., Roozendaal, B. & McGaugh, J. L. Glucocorticoid effects on object recognition memory require training-associated emotional arousal. Proceedings of the National Academy of Sciences 101, 853–858 (2004).

67. Vazdarjanova, A., McNaughton, B. L., Barnes, C. A., Worley, P. F. & Guzowski, J. F. Experience-dependent coincident expression of the effector immediate-early genes Arc and Homer 1a in hippocampal and neocortical neuronal networks. Journal of Neuroscience 22, 10067–10071 (2002).

68. Paxinos, G. & Franklin, K. B. The Mouse Brain in Stereotaxic Coordinates. (Gulf Professional Publishing, 2004).

69. West, M. J. Stereological methods for estimating the total number of neurons and synapses: Issues of precision and bias. Trends in Neurosciences vol. 22 51–61 (1999).

